# *Borrelia burgdorferi* BB0346 is an Essential, Structurally Variant LolA Homolog that is Primarily Required for Homeostatic Localization of Periplasmic Lipoproteins

**DOI:** 10.1101/2024.08.06.606844

**Authors:** Bryan T. Murphy, Jacob J. Wiepen, Danielle E. Graham, Selene K. Swanson, Maithri M. Kashipathy, Anne Cooper, Kevin P. Battaile, David K. Johnson, Laurence Florens, Jon S. Blevins, Scott Lovell, Wolfram R. Zückert

## Abstract

In diderm bacteria, the Lol pathway canonically mediates the periplasmic transport of lipoproteins from the inner membrane (IM) to the outer membrane (OM) and therefore plays an essential role in bacterial envelope homeostasis. After extrusion of modified lipoproteins from the IM via the LolCDE complex, the periplasmic chaperone LolA carries lipoproteins through the periplasm and transfers them to the OM lipoprotein insertase LolB, itself a lipoprotein with a LolA-like fold. Yet, LolB homologs appear restricted to γ-proteobacteria and are missing from spirochetes like the tick-borne Lyme disease pathogen *Borrelia burgdorferi*, suggesting a different hand-off mechanism at the OM. Here, we solved the crystal structure of the *B. burgdorferi* LolA homolog BB0346 (LolA_Bb_) at 1.9 Å resolution. We identified multiple structural deviations in comparative analyses to other solved LolA structures, particularly a unique LolB-like protruding loop domain. LolA_Bb_ failed to complement an *Escherichia coli lolA* knockout, even after codon optimization, signal I peptide adaptation, and a C-terminal chimerization which had allowed for complementation with an α-proteobacterial LolA. Analysis of a conditional *B. burgdorferi lolA* knockout strain indicated that LolA_Bb_ was essential for growth. Intriguingly, protein localization assays indicated that initial depletion of LolA_Bb_ led to an emerging mislocalization of both IM and periplasmic OM lipoproteins, but not surface lipoproteins. Together, these findings further support the presence of two separate primary secretion pathways for periplasmic and surface OM lipoproteins in *B. burgdorferi* and suggest that the distinct structural features of LolA_Bb_ allow it to function in a unique LolB-deficient lipoprotein sorting system.

**SIGNIFICANCE:** *Borrelia* spirochetes causing Lyme disease and relapsing fever have unusual double-membrane envelopes that instead of lipopolysaccharide (LPS) display abundant surface lipoproteins. We recently showed that secretion of these surface lipoproteins in *Borrelia burgdorferi* depends on a distant homolog of the canonical LPS outer membrane translocase LptD. Here, we probed the role of the *B. burgdorferi* Lol pathway in lipoprotein sorting and secretion. We show that the periplasmic chaperone LolA is essential, functionally different from *E. coli* LolA, with structural features of a bifunctional lipoprotein carrier protein operating without a downstream LolB outer membrane lipoprotein insertase. Depletion of LolA did not impact surface lipoprotein localization but led to a marked mislocalization of inner membrane lipoproteins to the outer membrane. This further supports two parallel, yet potentially interacting *Borrelia* lipoprotein transport pathways that are responsible for either secreting surface lipoprotein virulence factors or maintaining proper distribution of lipoproteins within the periplasmic space.

## INTRODUCTION

The Lyme disease spirochete *Borrelia burgdorferi* is a diderm bacterium with several envelope features that distinguishes it from other evolutionarily distant gram-negative model organisms, such as lacking lipopolysaccharide (LPS) on its outer surface, sequestering its flagella within the periplasm, or exhibiting only a relatively thin layer of peptidoglycan that is more closely associated with the inner membrane (IM) than the outer membrane (OM) of the cell (reviewed in ref. (1)). In lieu of LPS, *B. burgdorferi* expresses an arsenal of more than 90 surface lipoproteins, which are known to mediate the majority of identified interactions with tick vectors and mammalian hosts (reviewed in refs. (2, 3)) and appear to be secreted by a distantly related version of the Lpt LPS transport system (4). Eight subsurface lipoproteins have been identified at the inner leaflet of the OM (5). Currently, four of these subsurface lipoproteins have known functional roles in envelope biogenesis or pathogenesis: Lp6.6 (BBA62) is abundant in OM protein complexes, non-essential for in vitro growth but associated with the ability of the spirochetes to colonize ticks (6–9), BB0323 is involved in maintaining OM integrity that also affects transmission and pathogenicity (10–12), and BamD (BB0324) and BamB (BB0028) are the only two identified β-barrel assembly machinery (BAM) complex lipoproteins with discreet functions (13).

Proper localization of OM lipoproteins is critical to their function and first requires the export of pre-pro-lipoproteins with an N-terminal signal II sequence and lipobox motif through the cytoplasmic membrane via the general secretory (Sec) pathway. Following translocation, a conserved cysteine residue at the lipobox motif +1 site is modified by a series of three enzymes (Lgt, Lsp, and Lnt) to remove the signal sequence and generate mature tri-acylated lipoproteins at the periplasmic leaflet of the IM (reviewed in (14)). Extrusion of mature lipoproteins from the IM by the type VII ABC-transporter LolCDE ultimately determines which lipoproteins will finally be transported to the inner leaflet of the OM by the LolA periplasmic chaperone (15). In *E. coli* and other well studied γ-proteobacteria, intrinsic +2/+3/+4 “Lol sorting signal” residues immediately following the lipidated cysteine putatively interact with the primary membrane phospholipid phosphatidylethanolamine (PE) to retain IM lipoproteins. More specifically, anionic amino acids at these positions are thought to act as a Lol avoidance signal by interacting with cationic PE head amine groups (16). The additional clustered phospholipid chains from this interaction then prevent LolE from acquiring the lipoprotein and starting the extrusion process. While *Borrelia* has similarly charged phosphatidylcholine (PC) in place of PE, PC’s head amine group is sterically hindered by trimethylation (17). This may explain why no “+2/3/4” Lol sorting signals have been identified (5, 18, 19). Spirochetes, like α-, 8- and χ-proteobacteria, also lack a detectable homolog for the OM lipoprotein acceptor and insertase LolB, which acquires lipoproteins from LolA and inserts them into the periplasmic leaflet of the OM via a small protruding loop domain containing a functionally relevant leucine residue (20). Therefore, *Borrelia* likely uses different mechanisms for localizing periplasmic lipoproteins to the IM or the periplasmic leaflet of the OM.

Here, we began studying the putative LolB-deficient lipoprotein sorting system of *Borrelia burgdorferi* by solving the crystal structure of its LolA homolog BB0346 (*i.e.* LolA_Bb_), the furthest identifiable downstream component of the Lol pathway. We also tested for heterologous complementation of a conditional *E. coli lolA* knockout and used a conditional *B. burgdorferi lolA* knockout to assess the effects of LolA depletion on lipoprotein sorting and secretion. Our results provide first insights into the role of a structurally variant spirochetal LolA lipoprotein carrier protein homolog in maintaining a diderm envelope that is dominated by surface-localized lipoproteins.

## RESULTS

### *B. burgdorferi* BB0346 is a LolA homolog that fails to complement *E. coli* LolA

The NCBI Conserved Domain Database annotates the product of open reading frame (ORF) BB_0346, located on the linear chromosome of *B. burgdorferi*, as a LolA domain-containing protein (cd16325), which should function in the periplasmic space after transport across the inner membrane by SecYEG and subsequent cleavage by signal peptidase I (SPase I). Indeed, BB_0346 encodes for a 216-amino acid protein with a predicted 17 amino acid-long signal I peptide (SSI; SignalP 6.0 SSI probability = 0.97 (21)).

Initial heterologous expression of the full-length BB_0346 ORF, but not BB_0346 missing the SSI peptide, proved toxic to *E. coli*, inhibiting growth even from leaky expression under an uninduced *lac* promoter (data not shown). We hypothesized that full-length BB_0346 was toxic to *E. coli* because it either (i) was not properly recognized, processed, and secreted by the *E. coli* general secretion machinery, or (ii) was able to partially interfere with the periplasmic Lol system machinery in *E. coli*, leading to aberrant OM lipoprotein sorting. A comparison of the two signal I peptides showed that the *B. burgdorferi* SSI peptide was shorter and had a more degenerate signal I peptidase recognition sequence than that of *E. coli* (**Fig. 1A**). In a series of experiments, we generated various chimeric *E. coli/B. burgdorferi* LolA fusion proteins and tested them for toxicity and proper processing. Only the generation of a chimeric LolA construct fusing the *E. coli* LolA SSI peptide to the mature *B. burgdorferi* LolA (mLolA_Bb_) peptide eliminated toxicity and produced a major protein band that was equal in size to the native LolA_Bb_ produced in *B. burgdorferi* (**Fig. 1B**). This indicated that the *E. coli* SPase I (LepB) does not efficiently process the w.t. *B. burgdorferi* protein, but that processing efficiency is increased for an N-terminal chimera with the *E. coli* LolA signal peptide. Note that proper processing in *B. burgdorferi* appears independent of which SSI was used, suggesting that processing of secreted non-lipidated proteins is more promiscuous in that system. Fusions of either SSI to a superfolder GFP (sfGFP) reporter further demonstrated that the *B. burgdorferi* SSI interferes with processing and secretion of proteins into the *E. coli* periplasmic space, producing aggregates primarily located at the cell poles. In contrast, *E. coli* SSI-sfGFP fusions appeared more soluble and evenly distributed (**Supplemental Fig. S1**).

**Fig. 1.**
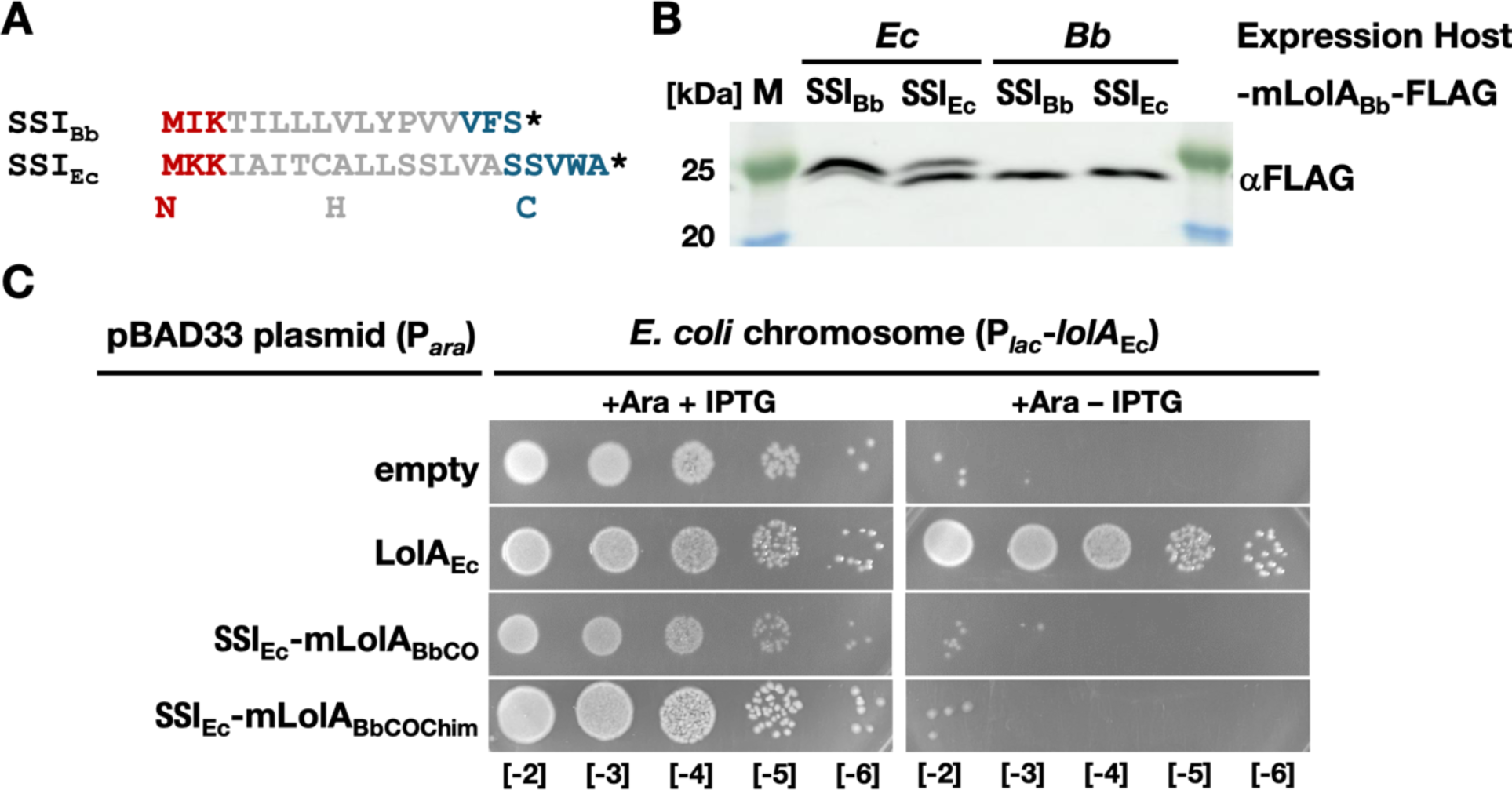
Complementation assay showing incompatibility of LolA_Bb_ with the *E. coli* Lol pathway. **(A)** Comparison of LolA secretory signal I peptides (SSI) in *B. burgdorferi* (Bb) and *E. coli* (Ec). The three regions of SSI peptides, including a positively charged N terminus (N), hydrophobic α-helix (H), and flexible C region (C), are denoted before the predicted signal peptidase I processing site (*). **(B)** Processing of heterologous LolA SSIs in *E. coli* and *B. burgdorferi.* C-terminally FLAG-tagged mature LolA_Bb_ (mLolA_Bb_-FLAG) fused to either *E. coli* or *B. burgdorferi* SSIs were overexpressed from P*_lac_* promoters in *E. coli* or *B. burgdorferi,* and whole cell lysates were analyzed by immunoblotting with anti-FLAG antibody. Note that the SSI_Bb_-mLolA_Bb_-FLAG fusion corresponds to a C-terminally FLAG-tagged w.t. LolA_Bb_ protein, marking the size of the properly processed LolA_Bb_. **(C)** Serial dilution spot plating of *E. coli* strain TT011 (P*_lac_*-*lolA*; (22)) harboring pBAD33 plasmid derivatives expressing various LolA constructs from the P*_ara_* promoter on LB agar containing 0.2% arabinose (Ara) with (+) or without (-) 1mM IPTG. SSI_Ec_-mLolA_BbCO_ includes mLolA_Bb_ that is codon-optimized for *E. coli*. SSI_Ec_-mLolA_BbCOChim_ is the C-terminal chimera based on the *C. vibrioides/E. coli* LolA chimera described by Smith and colleagues (23); see also **Supplemental Data**). Strains containing empty pBAD33 vector and recombinant *E. coli* LolA (LolA_Ec_)-expressing plasmids were included as negative and positive controls. Expression of the non-complementing LolA proteins was ascertained by Western immunoblotting (**Supplemental Data Fig. S1A**)

Building on these data, we next tested whether *B. burgdorferi* LolA was able to complement *E. coli* LolA using a conditional *E. coli* LolA knockout strain TT011 (22) (supplied by H. Tokuda). To reduce other potential issues for proper expression in *E. coli*, we further modified the apparently processable SSI_Ec_-mLolA_Bb_ fusion by codon optimization. We also generated an additional chimera that included a C-terminal adaptation which allowed a LolA homolog from the α-proteobacterium *Caulobacter vibrioides* to be compatible with the LolCDE complex in *E. coli* (23) (**Supplemental Data Fig. S2**). None of the chimeras were able to complement *E. coli* LolA and support growth (**Fig. 1C**), suggesting that *B. burgdorferi* LolA had functionally unique properties. It should be noted that *E. coli* TT011 still expresses OM lipoprotein Lpp, which is known to be toxic when mislocalized due to its linkage with peptidoglycan (24). As previously described, the *lpp-*minus TT011 derivative, TT015, showed no basic growth defect when LolA was depleted by removal of IPTG (22), so we were unable to test for complementation in that background.

### BB0346 (LolA_Bb_) is a structural LolA homolog with unique structural features

Based on the observed incompatibility with the *E. coli* Lol system, we wondered if the LolA_Bb_ structure might deviate from canonical LolA homologs. We therefore solved its structure by X-ray crystallography. A signal peptide-less BB_0346 sequence corresponding to the fully processed mature mLolA_Bb_ peptide was amplified by PCR from *B. burgdorferi* type strain B31 genomic DNA and ligated into pET29b(+), which provided a C-terminal hexa-histidine tag (**Supplemental Data Fig. S3A**). Note that the peptide sequences of BB0346/LolA_Bb_ from strains 297 and B31 are identical. The resulting plasmid was used to transform *E. coli* BL21(DE3) pLysS (Novagen). Soluble mLolA_Bb_ was purified from the bacterial cytoplasm via two rounds of cobalt affinity chromatography separated by a single round of ion-exchange chromatography (**Supplemental Data Fig. S3B**). Crystallization screening on the purified LolA_Bb_ is described in the Materials & Methods section, and representative crystals are shown in **Supplemental Data Fig. S3C**.

The final LolA_Bb_ model resolved to 1.9 Å and included residues spanning Q2 to Y195 (**Fig. 2**). The residues that could not be modeled due to disorder are noted in **Supplemental Data Fig. S3A**. The overall structure of LolA_Bb_ contains 12 β-strands and two α-helices. These secondary structure elements adopt a partial β-barrel fold composed of 11 antiparallel β-strands (β1-11) and a single short strand (β12) which are capped on one end by the two α-helices (**Figs. 2A-C**). Prominent positive electron density was observed within the interior of the β-barrel that was ultimately modeled as a PEG molecule fragment obtained from PEG 5000 MME in the crystallization solution (**Fig. 2D**). Therefore, this structure is referred to as LolA_Bb_-PEG (PDB accession number 7TPM) from this point forward. Data sets that were obtained with crystals obtained in the absence of PEG also showed electron density in the LolA_Bb_ core, albeit to a lesser extent. For example, refinement of LolA_Bb_ models against X-ray diffraction data collected on crystals obtained from Proplex HT G2 (2 M ammonium sulfate, 100 mM sodium acetate pH 5.0) and H1 (2 M NaCl, 100 mM sodium citrate pH 6.0) produced weaker electron density in the core of LolA_Bb_ (**Supplemental Data Fig. S4**). As such, it is conceivable that a polar lipid-like molecule was acquired from the *E. coli* expression host and bound to LolA_Bb_ at low occupancy. Additionally, we observed more prominent electron density in this region from crystals that were cryoprotected with PEG 200 instead of glycerol, suggesting that the PEG molecule may displace any molecules that are acquired from the expression host under these conditions (data not shown).

**Fig. 2.**
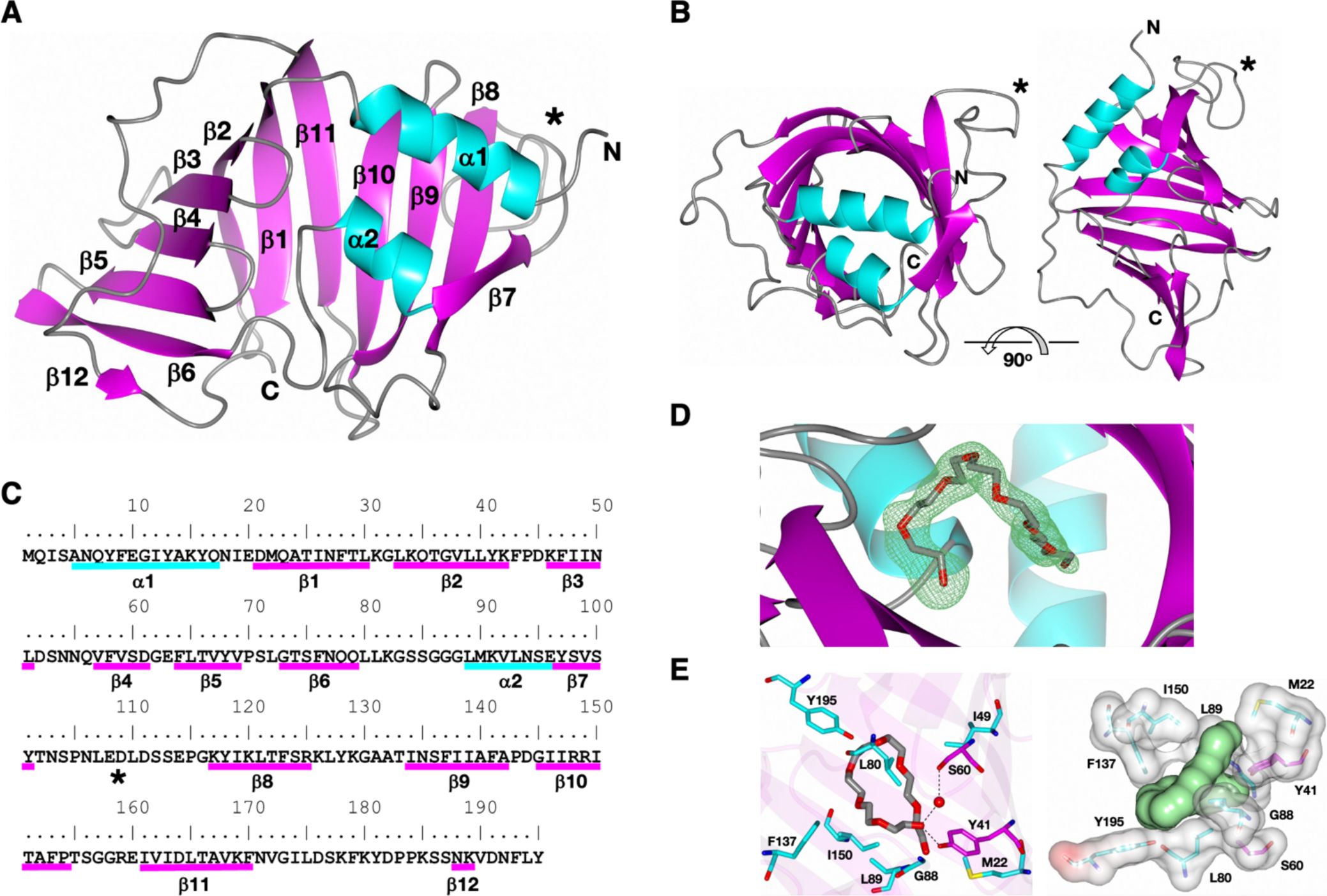
X-ray crystal structure of LolA_Bb_-PEG. **(A)** Ribbons rendering colored by secondary structure with α-helices (cyan) and β-sheets (magenta) indicated. N, N terminus; C, C terminus. An asterisk (*) indicates the location of a unique protruding loop between β7 and β8. **(B)** View of LolA_Bb_-PEG along and perpendicular to the β-barrel axis. **(C)** Secondary structure elements are annotated relative to the mLolA_Bb_-his sequence excluding C-terminal purification tag residues (see **Supplemental Data Fig. S3A**). **(D)** Fo-Fc omit electron density map (green mesh) contoured at 3σ showing the PEG molecule bound to LolA_Bb_. **©** Interaction of PEG with LolA_Bb_-PEG. In the left panel, the PEG molecule (gray/red cylinders) is shown to be surrounded mainly by hydrophobic residues (cyan). The direct hydrogen bond to Y41 and water mediated contact with S60 is indicated by the dashed lines. The right panel shows an electrostatic surface representation of the LolA_Bb_-PEG residues surrounding the PEG molecule (green).

Within LolA_Bb_-PEG, the PEG molecule is mainly surrounded by hydrophobic residues, but it forms a direct hydrogen bond to Y41 (OH) and water mediated contact with S60 (OH) as shown in **Fig. 2E**. It has a total accessible surface area of 574.4 Å^2^, of which 567.1 Å^2^ (98.7%) is buried in the core, forming an interface with LolA_Bb_-PEG that covers 461.9 Å^2^ as determined by PISA (25). As such, the PEG molecule is positioned within a hydrophobic pocket (**Fig. 2E**). In addition, a sodium ion was modeled based on coordination geometry and bond distances (**Supplemental Data Fig. S5**). Interestingly, subsequent LolA_Bb_ preparations also yielded crystals from 1.5 M ammonium sulfate, 0.1 M HEPES, pH 7.5 which were cryoprotected with 2.5M lithium sulfate (see Materials & Methods). In these crystals, a large mass of positive electron density within the PEG binding site was ultimately modeled as steric acid (LolA_Bb_-SA). The steric acid molecule adopted a binding mode similar to the PEG molecule, as shown in **Supplemental Data Fig. S6A**.

Superimposition of various solved LolA structures onto LolA_Bb_-PEG using secondary structure matching (SSM) (26) yielded the following RMSD deviations between Cα atoms: *Pseudomonas aeruginosa* (2W7Q, 2.86 Å, 146 residues), *E. coli* (1IWL, 2.95 Å, 131 residues), *Neisseria europaea* (3BUU, 2.50 Å, 137 residues), *Yersinia pestis* (4KI3, 2.50 Å, 130 residues). As shown in **Fig. 3**, there is displacement of both β-sheets and α-helices between these similar structures, and this accounts for the somewhat large RMSD deviations. One major difference is observed in the loop between β6 and α2 in LolA_Bb_-PEG (α3 in other LolA structures) which is moved out of the core to accommodate binding of the PEG molecule. In the γ-proteobacterial homologs, this region contains either an additional α-helix that is known to have important function in the transfer of lipoprotein cargo (27), or flexible loops that would both clash with the PEG molecule. Finally, superimposition of LolA_Bb_-SA and LolA_Bb_-PEG and the recently solved liganded *E. coli* LolA-Pal_13_ lipopeptide complex (28) (7Z6W) yielded an RMSD of 2.44 Å between Cα atoms (136 residues) and confirmed that the PEG and SA molecules bind at the site that is functionally occupied in *E. coli* LolA by the Pal_13_ lipopeptide (**Supplemental materials Fig. S6B**). Intriguingly, *E. coli* LolB (1IWN) also has a reported PEG fragment bound to the core of the protein. Superimposition of LolB with LolA_Bb_ yielded an RMSD deviation of 2.84 Å for 127 aligned residues. As shown in **Fig. 4**, the structures are similar, but they contain displaced secondary structure elements relative to one another, including an additional α-helix that would clash with the PEG molecule in LolA_Bb_, as already noted for some of the γ-proteobacterial LolAs. Notably, the PEG molecules in LolB and LolA_Bb_-PEG adopt similar binding modes (**Fig. 4B**). However, the PEG molecule in LolA_Bb_-PEG is a larger fragment which requires displacement of the loop between β6 and α2 for binding.

**Fig. 3.**
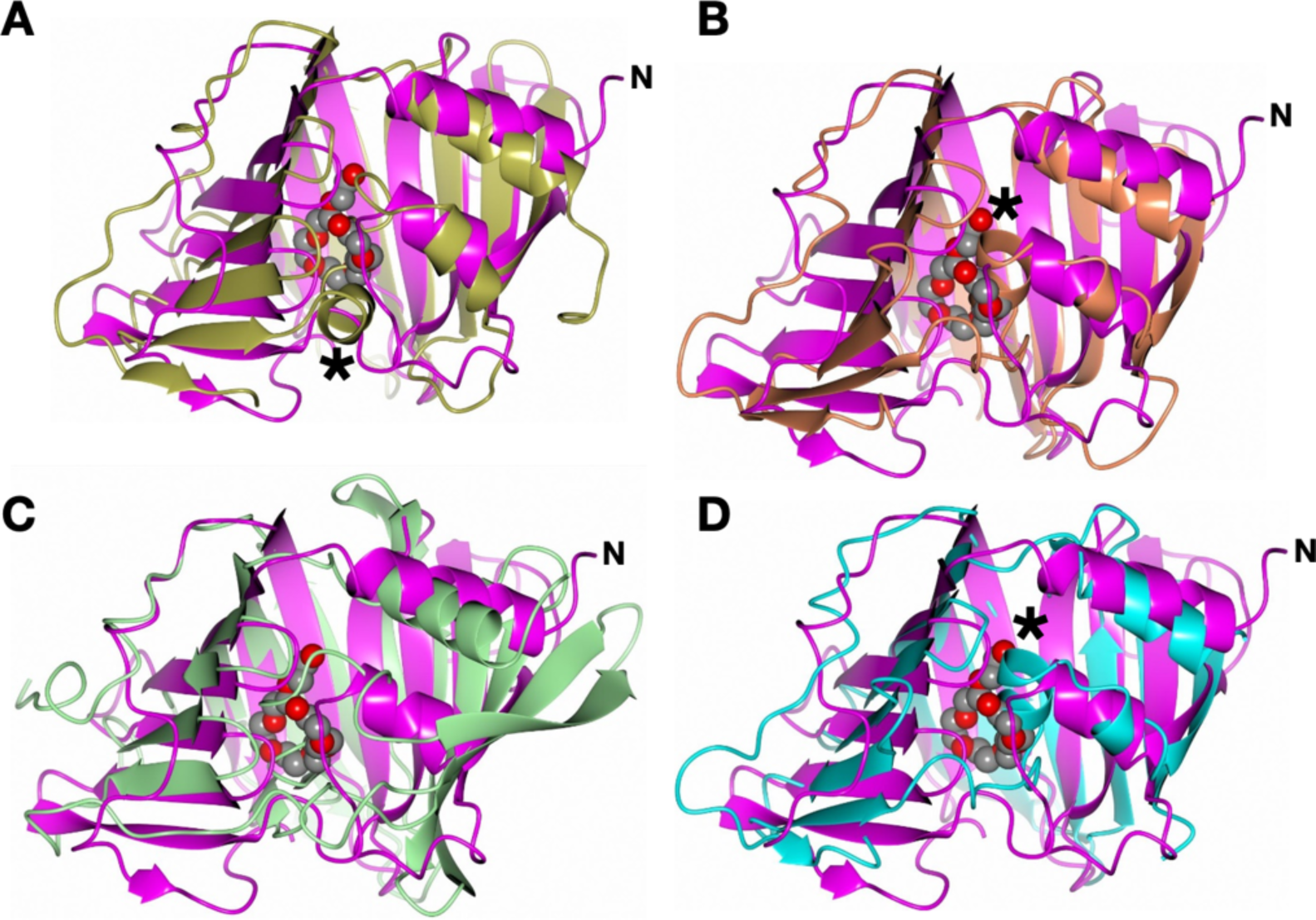
Structural differences between the hydrophobic pockets of LolA homologs. LolA_Bb_-PEG was superimposed with the LolA structures of **(A)** *P. aeruginosa* (2W7Q, gold), **(B)** *E. coli* (1IWL, coral), **(C)** *N. europaea* (3BUU, green) and **(D)** *Y. pestis* (4KI3, cyan) onto LolA_Bb_ (magenta). The PEG molecule from LolA_Bb_ is rendered as gray/red spheres. The additional α-helix present in the other LolA structures is indicated by asterisks.

**Fig. 4.**
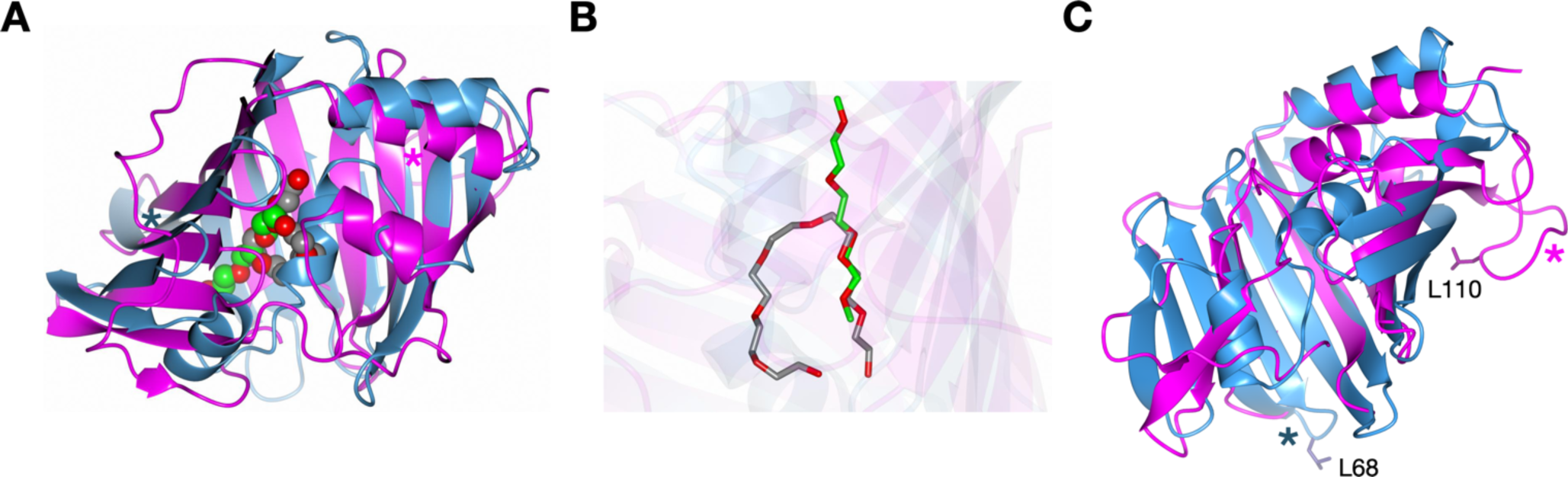
Structural comparison of LolA_Bb_ and *E. coli* LolB. **(A)** The structure of *E. coli* LolA (1IWN, blue) was superposed onto LolA_Bb_-PEG (magenta). The PEG molecule is rendered in gray/red spheres for LolA_Bb_-PEG and in green/red spheres for LolB. Localization of the two protruding loops in LolA_Bb_ and *E. coli* LolB are indicated with colored asterisks. **(B)** Binding modes for the PEG molecules bound to LolB (green/red) and LolA_Bb_ (gray/red). **(C)** Rotation of superimposed structures shown in panel A to better illustrate the localization of the two loops, marked with colored asterisks as in panel A. The functionally relevant *E. coli* LolB L68 residue and a similarly prominent LolA_Bb_ L110 residue of are shown as sticks.

In addition to the absence of an α-helix, rotation of the four structure overlays shown in **Fig. 3** reveals another significant difference between LolA_Bb_-PEG and other solved homologs (**Fig. 5)**. More specifically, LolA_Bb_-PEG has a 15 amino acid flexible loop spanning residues T102 to G116 that replaces a turn between antiparallel β-sheets 7 and 8 in the γ-proteobacterial LolA homologs. This solvent-exposed flexible loop contains two hydrophobic leucine residues (L107 and L110) near the loop’s center point with predicted salt bridging from D111 to the backbone’s R148 and R159 residues. Together, L110, D111, R148, and R149 compose 4 of the most highly conserved residues among 150 diverse spirochetal homologs including those found within Lyme disease and relapsing fever-causing *Borreliaceae,* syphilis and periodontal disease-causing *Treponemataceae*, and other environmental spirochetes (**Supplemental Data Fig. S7)**. Intriguingly, LolB from *E. coli* has a similar hydrophobic residue-containing loop that is responsible for inserting lipoproteins into the periplasmic leaflet of the OM with its central L68 residue (20). However, the LolB loop is composed of only five amino acids and is located at the opposite end of LolB when the structures are superimposed **(Fig. 4).**

**Fig. 5.**
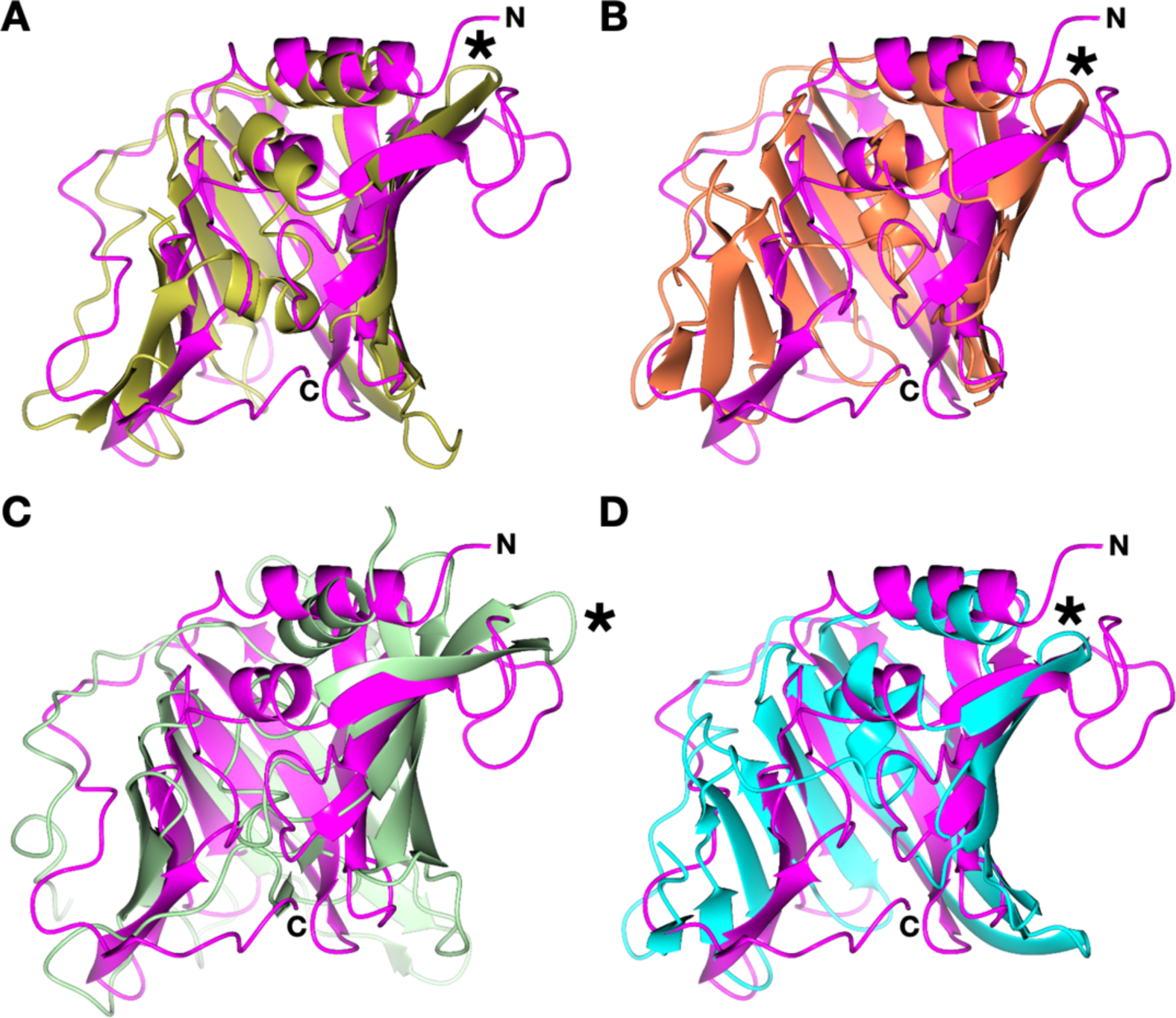
Structural differences between LolA homologs in the β7-β8 connecting loop. LolA_Bb_-PEG was superimposed with the LolA structures of **(A)** *P. aeruginosa* (2W7Q, gold), **(B)** *E. coli* (1IWL, coral), **(C)** *N. europaea* (3BUU, green) and **(D)** *Y. pestis* (4KI3, cyan) onto LolA_Bb_ (magenta). The turn between antiparallel β7 and β8 sheets present in the other LolA structures is indicated by asterisks.

### LolA_Bb_ is essential for *B. burgdorferi* growth

To investigate whether LolA_Bb_ is required for growth, we used a conditional *lolA* knockout strain of *B. burgdorferi* 297. In this recombinant strain, a kanamycin resistance cassette was used to disrupt the chromosomal copy of BB_0346 while expressing an ectopic plasmid-encoded allele under an IPTG-inducible P_pQE30_ promoter. As shown in **Fig. 6A**, removal of IPTG driving the expression LolA_Bb_ led to a marked growth defect within 48 hours. By day 2 post-IPTG removal, we observed pleomorphic abnormalities that ranged from blebbing along the cell length to full rounding of some cells, suggesting severe structural deficiencies in the cell envelope (**Fig. 6B**). At day 1 post-depletion, we detected an approximate 10-fold reduction in LolA_Bb_ protein levels compared to w.t. cells **(Fig. 6D)**. This was concomitant with a 12-fold reduction in BB_0346 transcript levels, as measured by RT-qPCR (data not shown).

**Fig. 6.**
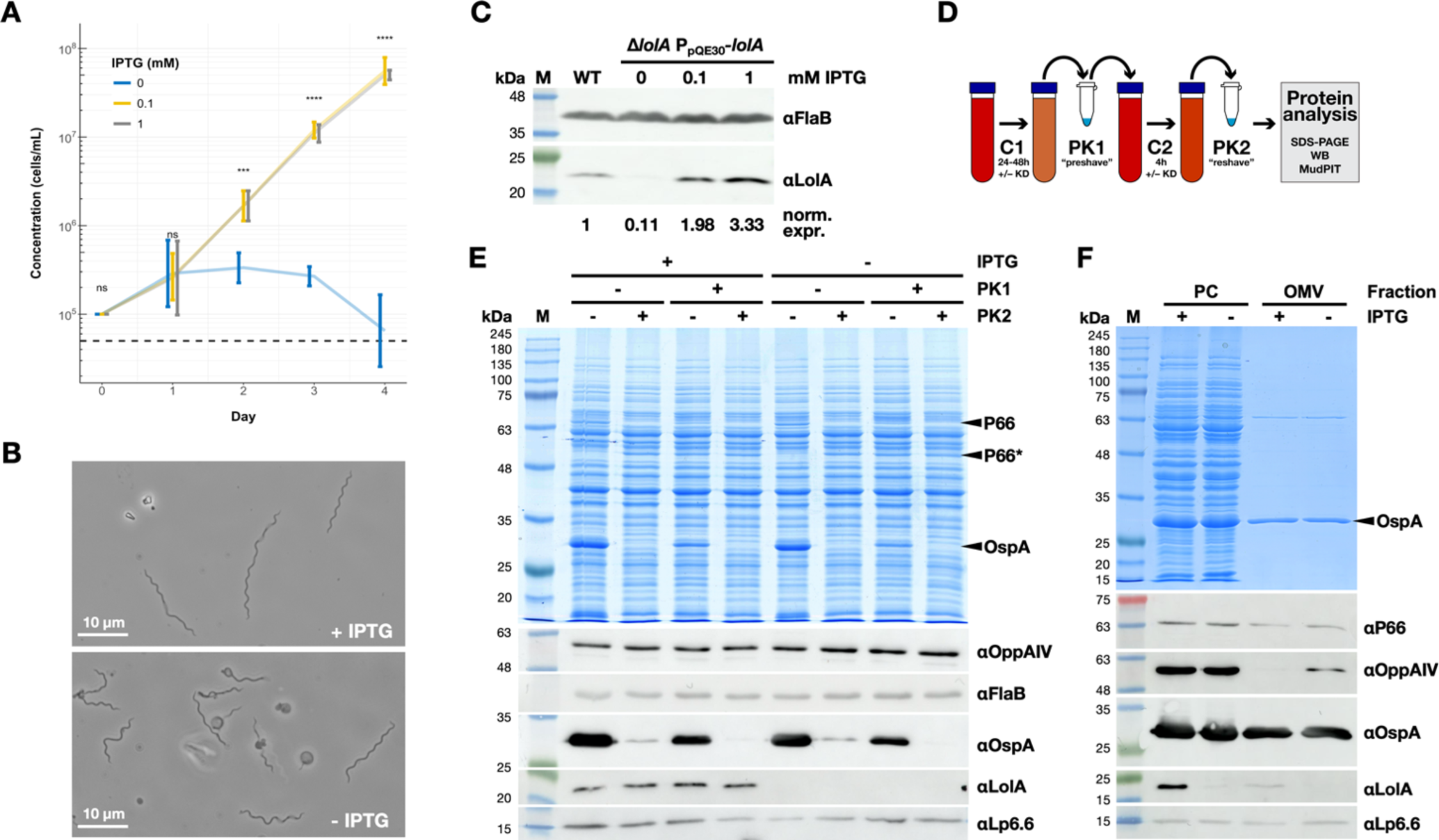
Phenotypic analysis of LolA_Bb_ depletion in *B. burgdorferi.* **(A)** Liquid culture growth curves. Spirochetes were inoculated at 1 × 10^5^ cells/mL and grown with or without IPTG to control expression of LolA_Bb_ in a recombinant *B. burgdorferi* strain carrying its sole plasmid-encoded *lolA*_Bb_ allele under *lac* promoter control (see Materials & Methods). Growth curves are from 3 biological replicates. Error bars indicate mean ± 95% confidence interval. Significance was calculated by 2-way ANOVA; \*\*\**P* ≤ 0.001; \*\*\*\**P* ≤ 0.0001; NS, *P* > 0.05. **(B)** Phase contrast micrographs depicting the effects of LolA_Bb_ depletion (-IPTG) on cell morphology in comparison to control cells (+ IPTG) on day 2. **(C)** Expression levels of LolA_Bb_ in w.t. and recombinant conditional knockdown strains. Periplasmic flagellar protein FlaB served as a loading control. Expression levels relative to w.t. cells were determined by densitometry measurements of Western immunoblot bands from 3 biological replicates. **(D)** Schematic depiction of modified surface protein accessibility assay (see text). **(E)** Surface accessibility of lipoproteins using a modified PK shaving/reshaving assay (see panel D), as analyzed by Commassie-stained SDS-PAGE, Coomassie staining and Western immunoblotting. Highly abundant OspA served as the model surface lipoprotein. OM porin P66 served as a non-lipoprotein OM control; note that after PK cleavage of a surface-accessible loop, a P66 fragment band (P66*) appears. Periplasmic flagellar protein FlaB was used as an OM integrity and constitutively expressed loading control. OppAIV and Lp6.6 served as sentinel inner membrane and subsurface lipoprotein controls, respectively. **(F)** Fractionated protoplasmic cylinder (PC) and outer membrane vesicle (OMV) fractions were assessed as in **panel E**. Note that the PC fraction also contains intact cells.

### LolA_Bb_ depletion does not affect surface lipoprotein secretion but leads to emerging mislocalization of IM lipoproteins to the OM

The observed delay between LolA_Bb_ depletion and phenotypic changes was likely a consequence of envelope homeostasis mediated by residual, properly localized lipoproteins. To observe nascent surface lipoprotein secretion under LolA_Bb_-depleted conditions, we modified our standard proteinase K (PK) surface localization assay (29) by adding a “pre-shaving” step. Briefly, cells cultured under LolA_Bb_-depleting and control conditions for 24 hours (at the day 1 timepoint) were washed and treated with PK for 1 hour to remove accessible surface-localized proteins. After a 4-hour recovery in fresh culture medium under depleting or non-depleting conditions, the cells were again washed and then “re-shaved” with PK (**Fig. 6C**). Western immunoblot analysis of the resulting samples showed that LolA_Bb_-depleted cells maintained their ability to secrete the prototypical outer surface lipoprotein OspA, presumably using a reconfigured Lpt transport system in *Borrelia burgdorferi* (4). At the same time, periplasmic IM lipoprotein OppAIV and OM lipoprotein Lp6.6, as well as the periplasmic flagellar protein FlaB, remained inaccessible to PK, indicating that they remained periplasmic and that there was no major disruption of the OM barrier at this timepoint. (**Fig. 6E**).

To further characterize the broader consequences of LolA_Bb_ depletion on OM lipoprotein transport, we next used a cell fractionation assay to separate and purify outer membrane vesicles (OMVs) from a protoplasmic cylinder (PC) fraction (30); note that the PC fraction also contains remaining intact cells. As expected from the PK shaving assays, there was no apparent defect in OspA transport to the OM under LolA_Bb_-depleting conditions. To our surprise, however, there was also no detectable change in Lp6.6 abundance in the OM, but OppAIV became significantly mislocalized to the OM (**Fig. 6F**).

To exclude the possibility of OMV contamination by IM proteins, and at the same time gain a more comprehensive view of the effect of LolA_Bb_ depletion on the OM proteome, we analyzed the PC and OMV cell fractions by Multidimensional Protein Identification Technology (MudPIT) label free quantitative mass spectrometry, as used in our previous studies of the *B. burgdorferi* envelope (4, 5). Based on the obtained <u>d</u>istributed <u>N</u>ormalized <u>S</u>pectral <u>A</u>bundance <u>F</u>actor (dNSAF) values, we were able to derive the abundance of each protein in each cell fraction. To calculate a dNSAF ratio illustrating abundance changes in the OM proteome, we then divided the OMV dNSAF values for each protein under LolA_Bb_-depleted conditions by those under control (“LolA_Bb_-replete”) conditions. Only proteins that were detected in at least 2 out of the 3 biological replicates were included in the analysis.

These data illustrated several points. First, they showed that the OMV fractions obtained on day 1 retained high OM-specific purity even with LolA_Bb_ depletion: non-lipoprotein IM controls such as secretory machinery proteins SecY, SecE, SecF and SecD, post-translational lipoprotein modification pathway proteins Lgt and Lnt, as well as LolD were readily detected in the PC fractions but remained undetectable in the OMV fractions (**Fig. 7**). Second, they further supported the intriguing OppAIV immunoblot results, as 6 of the 17 detected IM lipoproteins (including OppAIV) showed a statistically significant increase in OMV fraction abundance under LolA_Bb_ depletion conditions (**Fig. 8**), mostly above a mean 1.5-fold change. At the same time, only 3 of 24 detected surface lipoproteins were significantly reduced in the OMV fraction with LolA_Bb_ depletion, and their mean fold changes remained below 1.5-fold, confirming the OspA immunoblot data. Interestingly, the 6 detected periplasmic OM lipoproteins showed an almost even split between increased, decreased, and unchanged abundance: Lp6.6 and BB0323 increased significantly in abundance in the OMV fraction, whereas BamB and BB0460 dropped in abundance. Notably, there was no change in the OM abundance of BamD at this timepoint. All proteins with specific changes in abundance in the PC and OMV fractions are listed in **Supplemental Data Tables S2 and S3**, respectively.

**Fig. 7.**
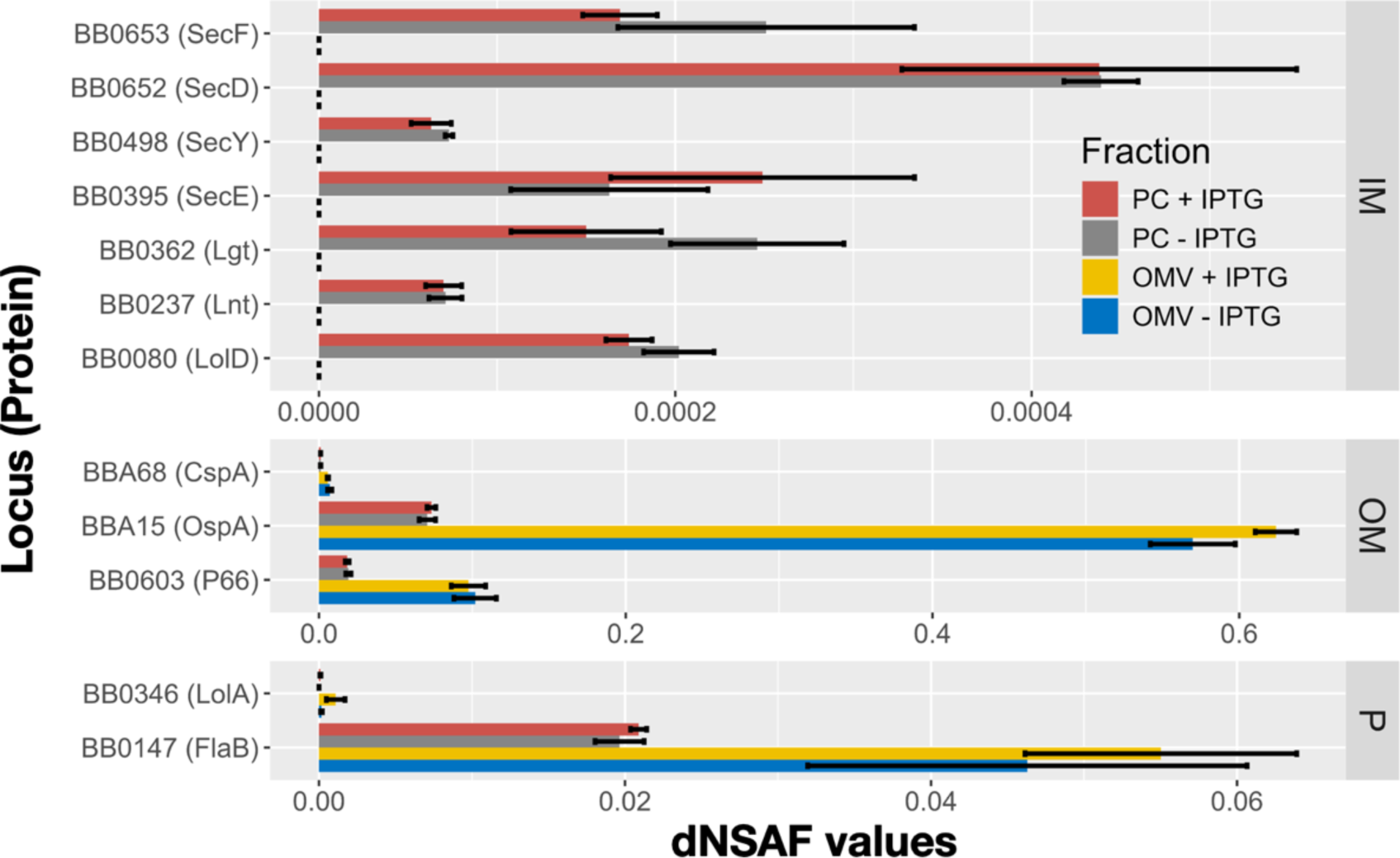
MudPIT analysis of *B. burgdorferi* conditional knockout strain cell fractions. dNSAF values indicating abundance of selected inner membrane (IM), outer membrane (OM) and periplasmic (P) proteins in the protoplasmic cylinder (PC) and outer membrane vesicle (OMV) fractions are plotted. + IPTG, LolA replete control conditions; - IPTG, LolA depleting conditions. Samples were taken at day 1 (24 hours depletion). Note that the PC fraction also contains remaining intact cells. Data are from 3 parallel biological replicates. Error bars indicate mean ± SEM.

**Fig. 8.**
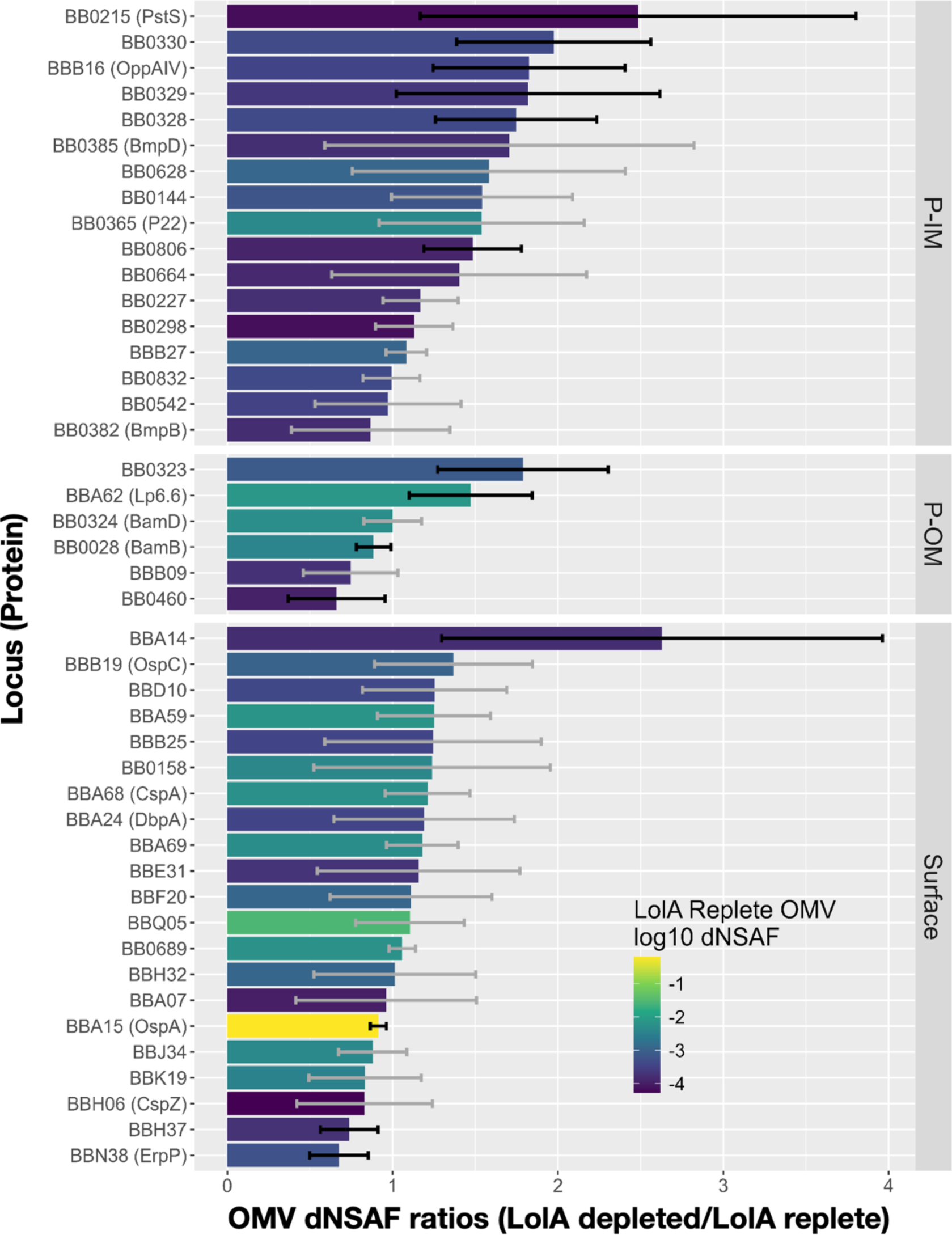
MudPIT analysis of *B. burgdorferi* conditional knockout strain cell fractions. dNSAF values indicating abundance of periplasmic inner membrane (P-IM), periplasmic outer membrane (P-OM) and surface lipoproteins consistently detected in the outer membrane vesicle (OMV) fraction were used to determine dNSAF ratios indicating a change in abundance. dNSAF ratios of LolA depleted (-IPTG) dNSAF value/LolA replete (+IPTG) are plotted. Samples were taken at day 1 (24 hours depletion). Data are from 3 parallel biological replicates. Error bars indicate mean ± SEM, wit propagation of uncertainty for ratio values. Column shading indicates the dNSAF-derived abundance of proteins under LolA replete control conditions; e.g., OspA (yellow) is much more abundant in the sample than BBA14 (purple).

To potentially demonstrate a more severe lipoprotein mislocalization phenotype, we attempted to characterize envelope fractions from cells that had been LolA_Bb_-depleted for 48 hours (day 2 timepoint). However, at this point, cell envelopes were apparently too disturbed (see **Fig. 6B**) to be successfully fractionated, resulting in OMV fractions that were indistinguishable in protein content from PC fractions when analyzed by SDS-PAGE (**Supplemental Data Fig. S8**).

## DISCUSSION

Efficient and accurate lipoprotein localization is essential for survival in diderm bacteria, as lipoproteins mediate a wide variety of important cellular functions, including the assembly and function of essential outer membrane machinery like the Bam complex (1). Like α-, δ-, and ε-proteobacteria, *B. burgdorferi* has all the canonical Lol pathway components found in *E. coli* except for an identified LolB outer membrane lipoprotein acceptor. Here, we solved the crystal structure of the *B. burgdorferi* LolA homolog BB0346 (LolA_Bb_) in both PEG and steric acid bound forms to 1.9 Å and 1.8 Å resolution, respectively. In these structures, we identified two unique LolA_Bb_ loop domains with comparative analyses to other solved LolA structures.

The first loop replaces an entire α-helix (α2) that typically protrudes into the hydrophobic core of canonical LolA homologs to facilitate cargo acceptance from LolC at the IM and transfer to LolB at the OM (27) (**Fig. 3**). This loop appears to provide increased flexibility over the α-helix, since it is fully displaced out of the protein’s core to accommodate binding of a PEG molecule fragment or steric acid. Curiously, AlphaFold (AF-O51321-F1) predicts this loop is also displaced out of the core in the absence of ligand, providing a fairly accurate prediction that differs from the solved LolA_Bb_-PEG structure (7TPM) by an RMSD of 1.077 Å. The second loop extends a turn between antiparallel β-sheets 7 and 8 near the N-terminus of LolA_Bb_ (**Fig. 5**), bearing some resemblance to a LolB loop that is involved in lipoprotein insertion into the outer membrane (20) (**Fig. 4**). Recently, Smith and colleagues identified a “bifunctional” LolA homolog (LolA_Cv_) from the also LolB-deficient α-proteobacterium *Caulobacter vibrioides* (23). LolA_Cv_ was modeled *in silico* to also have an extended loop between antiparallel β-sheets 7 and 8 (loopβ7-β8), and this loop was experimentally linked to the protein’s ability to insert lipoproteins into the OM by replacing LolA and LolB in *E. coli* with heterologously expressed LolA_Cv_ (23). Notably, deletion or substitution of L119 near the center point of loopβ7-β8 in LolA_Cv_ eliminated its ability to complement *E. coli* LolB. This suggests that the corresponding crystal structure-confirmed loopβ7-β8 in LolA_Bb_ provides a similar LolB-like lipoprotein insertase function. Of note, the positively charged C-terminus of LolA_Cv_ had to be modified to allow for interaction with the periplasmic pad of *E. coli* LolC (23). In this study, a similarly modified LolA_Bb_ chimera failed to complement *E. coli* LolA (**Fig. 1; Supplemental Fig. S2**), suggesting that the spirochetal LolA homolog might function differently still.

There are several potential explanations for why chimeric LolA_Bb_ was unable to complement the *E. coli* LolA conditional knockout strain despite its correct localization to the periplasmic space and C-terminal alteration for permitting interaction with the LolCDE complex in *E. coli*. For example, the replacement of the canonical LolA α2-helix by a flexible loop in *B. burgdorferi*, and potentially all *Borreliaceae* homologs, may lead to incompatibility between proteobacterial and spirochetal Lol systems in both the lipoprotein acceptance and release steps. Another possible explanation is rooted in the different phospholipid compositions of *E. coli* and *B. burgdorferi*. Just as phospholipid composition was shown to be important for extrusion of lipoproteins from the inner membrane by LolE (16), a soluble form of LolB (denoted mLolB) was able to insert lipoproteins into synthetic liposomes in a phospholipid composition-dependent manner (31). The particularly phosphatidylcholine-rich membranes of *Borrelia* may therefore be required for LolA_Bb_ insertase function. Finally, our experiments with a *B. burgdorferi* conditional knockout strain resulted in unexpected mislocalization of IM lipoproteins to the outer membrane under LolA_Bb_ depletion conditions (**Figs. 8 and 9**). One possible explanation for this phenotype might be that disruption of the Lol pathway leads to an accumulation of lipoproteins in the IM, which are then indiscriminately force-fed to the otherwise surface lipoprotein-transporting Lpt pathway to aleviate IM stress. As we didn’t observe sudden surface exposure of periplasmic OM lipoproteins under those conditions, this would require ejection of natively non-surface lipoproteins from the Lpt pathway at the periplasmic face of the OM. Another, even more intriguing explanation would be that the *B. burgdorferi* Lol pathway works not only in the canonical anterograde direction but can also remove mislocalized lipoproteins from the OM and return them to the IM in a retrograde step, reminiscent of the proteobacterial Mla (<u>m</u>aintenance of OM lipid <u>a</u>symmetry) system removing misplaced phospholipids from the surface leaflet of the proteobacterial OM and transporting them back to the IM (32, 33).

More detailed studies will be needed to investigate the operational directionality and precise structure-function of LolA_Bb_ in its native system, since our current observations suggest that the evolutionary distance between spirochetes and proteobacteria may have resulted in diverse Lol system modalities. Such was the case for the *B. burgdorferi* BB0838 homolog of LptD that, rather than transporting lipidated polysaccharides, was shown to be required for the translocation of lipidated proteins to the LPS-deficient cell surface of *Borrelia* (4). In support of this previous finding, we have now observed that LolA_Bb_ plays no direct role in the transport of surface lipoproteins, as depletion of LolA_Bb_, unlike depletion of LptD_Bb_, had little to no immediate effect on the proper localization of OspA and other surface lipoproteins. This further supports our earlier proposed model of two dichotomous lipoprotein pathways in *Borrelia* spirochetes (4): (i) a pathogenesis-associated Lpt pathway that ensures rapid deployment of crucial surface lipoprotein virulence factors required for efficient vector-borne transmission, dissemination, and persistent infection of vectors and reservoir hosts, and (ii) a “house-keeping” Lol pathway that ensures proper lipoprotein localization and homeostasis within the periplasm. Specific mechanistic details on both transport pathways have yet to be elucidated, including the possibility for some collaborative crosstalk between them. Since both systems are essential but composed from modular components that diverge structurally and functionally from those found in other model diderms, their continued study will further highlight the diversity of microbial envelope biogenesis systems and potentially lead to the discovery of narrow-spectrum therapeutics.

## MATERIALS & METHODS

### Strains and growth conditions

Chemically competent NEB^®^ 5-alpha F’*Iq* cells (NEB; C2992H) were transformed with recombinant plasmids per manufacturer instructions and grown at 37°C on selective Luria-Bertani (LB) agar plates (BD; 244520) and in LB broth (Fisher; BP1426). Plasmid DNA was isolated from *E. coli* clones using a Miniprep kit (Macherey-Nagel; 740588) and verified by Oxford Nanopore Technology sequencing (Plasmidsaurus) or single-pass primer extension sequencing (ACGT, Inc.) with DNA oligonucleotide primers (Integrated DNA Technologies). Verified plasmids were then used to transform *E. coli* or *B. burgdorferi* strains as specified. The *E. coli* TT011 conditional knockout of *lolA* was kindly provided by Hajime Tokuda ((22); Institute of Molecular and Cellular Biosciences, University of Tokyo, Tokyo, Japan).

*B. burgdorferi* strains B31 and 297 and their recombinant derivatives were grown at 34°C under 5% CO_2_ in sterile filtered Barbour-Stonner-Kelly-II (BSK-II) medium containing 9.7 g/L CMRL-1066 (US Biological, C5900-01), 5.0 g/L neopeptone (Gibco, 211681), 6.6 g/L HEPES sodium salt (Fisher, BP410), 0.7 g/L citric acid (Sigma, C-8532), 5.0 g/L dextrose anhydrous (Fisher, BP350), 2.0 g/L yeastolate (Gibco, 255772), 2.2 g/L sodium bicarbonate (Fisher, BP328), 0.8 g/L sodium pyruvate (Fisher, AC132155000), 0.4 g/L *N*-acetylglucosamine (Sigma, A3286), 25 mg/L phenol red (Sigma, P-3532), and 50.0 g/L bovine serum albumin (Gemini, 700-104P) at pH 7.6-7.7, with 60 mL/L heat-inactivated rabbit serum (Pel Freez, 31126), and 200 mL/L 7% gelatin (Gibco, 214340) added before use. Streptomycin was added to BSK-II at a final concentration of 100 µg/mL for selection of *B. burgdorferi* strains containing IPTG-inducible plasmids.

### Recombinant plasmids

Recombinant plasmids were produced by a combination of classical subcloning, gene splicing by overlap extension (SOEing), site-directed mutagenesis (SDM), and DNA fragment synthesis through Twist Bioscience. For plasmid maps and sequences, see **Supplemental Materials and Methods**). To simplify the cloning of genes under an IPTG-inducible P*_pQE30_* promoter for controlled expression in both *E. coli* and *B. burgdorferi*, we modified pJSB104 (34) by removing an extraneous NdeI site in the plasmid’s backbone via SDM with primers Nde-aadA_F (5’-GAGGTTTCCAGATGAGGGAAGCGGTGATC-3’) and Nde-aadA_R (5’-CTTCCCTCATCTGGAAACCTCCCTCATTTAAAATTG-3’). Fusions of either the *E. coli* or *B. burgdorferi* signal peptide sequence I (SSI) to superfolder GFP (sfGFP) or the mature region of BB0346 with a C-terminal FLAG tag (mLolA_Bb_-FLAG) were then restriction-ligated into the modified pJSB104, resulting in expression plasmids for SSI_Ec_-sfGFP, SSI_Bb_-sfGFP, SSI_Ec_-mLolA_Bb_-FLAG, or SSI_Bb_-mLolA_Bb_-FLAG. Spectinomycin was added to LB and LB agar at a final concentration of 100 µg/mL for the maintenance of all IPTG-inducible plasmids in *E. coli*.

pBAD33 (35) (provided by Joe Lutkenhaus, University of Kansas Medical Center) was used for arabinose-inducible expression of genes in the *lolA* conditional knockout *E. coli* strain. First, an untagged *E. coli* codon-optimized SSI_Ec_-mLolA_Bb_ was synthesized by and subcloned into pBAD33 using SacI/HindIII restriction and ligation. Next, an upstream Shine–Dalgarno sequence from bacteriophage T7 gene 10 (36) was inserted via site directed mutagenesis with phosphorylated primers pBAD_RBS_F (5’-TTAAGAAGGAGATCGAGCTCATGAAAAAAATAGC-3’) and pBAD_RBS_R (5’-AGTTAAACAAAATTATTTCTAGCCCAAAAAAACGGG-3’). This plasmid was then modified for SPI_Ec_-mLolA_BbCOChim_ chimera expression by inserting a synthesized fragment digested with BglII and HindIII into the downstream region. Finally, the LolA_Ec_ expression vector was generated by replacing SPI_Ec_-mLolA_BbCO_ with a synthesized *lolA* gene from *Escherichia coli* str. K-12 substr. MG1655 via SacI/HindIII subcloning. Chloramphenicol was added to LB and LB agar at a final concentration of 34 µg/mL for the maintenance of all arabinose-inducible plasmids in *E. coli*.

### Transformation and clonal selection of *B. burgdorferi*

Electrocompetent B31-e2 or 297 cells were transformed by electroporation (37) with 500 ng to 5 μg of plasmid in 2-mm gap cuvettes (Thermo Scientific; 5520) using a Bio-Rad MicroPulser on EC2 setting, consistently measuring 2.49-kV/cm field strength and approximately 5-ms pulse times. Electroporated cells were immediately resuspended in 12 mL of prewarmed BSK-II and allowed to recover at 34°C for 18 to 20 h. Clonal selection of transformants was carried out by adding the 12 mL recovered culture to 35 mL selective BSK-II, followed by plating into 96-well microtiter plates and 8 to 16 days of incubation (38). Culture-positive wells were expanded into 6 mL of selective BSK-II and allowed to reach stationary phase for verification of plasmid acquisition by direct PCR of cultured *B. burgdorferi* cells using QuickLoad 2× *Taq* Master Mix (NEB, M0271). Positive clones were flash frozen on dry ice in BSK-II containing 10% DMSO (Sigma; D2438) and stored at −80°C.

### Recombinant *B. burgdorferi* strains

The *B. burgdorferi* conditional *bb_0346*/*lolA_Bb_* knockout was generated using a merodiploid intermediate. Endogenous plasmid contents of all clones were confirmed by PCR-based plasmid profiling (39), and only clones with profiles comparable to the strain 297 parent were used (data not shown). The BB_0346 ORF was amplified by PCR with flanking NdeI and HindIII sites; 5’ BB0346-NdeI (5’-GAGTTGGACATATGATAAAAACAATAC-3’) and 3’ BB0346-HindIII 5’-CATTTTCTTTCATAGATTGGAAGCTTAATTTTTTTTAA-3’). An internal HindIII site in *bb_0346* was removed by silent mutation using PCR SOEing; 5’ BB0346 int-HindIII (5’-AACCTTTTCTAGAAAACTTTACAAGGG-3’) and 3’ BB0346 int-HindIII (5’-CCCTTGTAAAGTTTTCTAGAAAAGGTT-3’). This NdeI/HindIII fragment was then inserted into a pJSB104 (34) derivative conferring streptomycin resistance, resulting in piBB0346. piBB0346 was electroporated into *B. burgdorferi* strain 297, yielding recombinant strain FF2. Plasmid recovery in *E. coli* was used to confirm the presence of piBB0346 in streptomycin-resistant transformants. Next, a pGEM-T Easy-based plasmid was used to disrupt the chromosomal copy of *bb_0346*. Regions upstream and downstream of the ORF were amplified by PCR using primers 5’ F1 bb0346 (5’-GAATATAGGGTAAGATAATTGCTGCTCGGC-3’), 3’ F1 bb0346-AscI (5’-gGcGcGCcGCTACTATTAATTCTTTTATTATTGCTTTTGC-3’), 5’ F2 bb0346-AscI (5’-ggcgcGCcGATATTTGAGAAAACACAACAACAGG-3’), and 3’ F2 bb0346-BssHII (5’-gcgcgcTGCCCTTTTTATATGCTTTAAAATATTGCAAGGC-3’), TA cloned and ligated with an AscI-flanked *aph[3’]-IIIa* kanamycin resistance marker (40) at the junction of the two fragments. This resulted in replacement of an internal 382-bp region of the *bb_0346* with the *aph[3’]-IIIa* marker, leaving 59 bp of the 5’ end ORF and 210 bp of the 3’ end of the ORF. The resulting mutation construct, designated pGEM-*bb0346::aph[3’]-IIIa*, was used to transform recombinant strain FF2. Transformants were selected with streptomycin/kanamycin and grown in BSK-II supplemented with 1 mM IPTG to maintain *bb_0346* expression. The presence of piBB0346 in two kanamycin/streptomycin-resistant clones, FH2 and FH4, was confirmed by plasmid recovery, and PCR was used to confirm disruption of the chromosomal copy of *bb_0346* by the *aph[3’]-IIIa* marker. The resulting chromosomal locus is shown in **Supplemental Materials Fig. S10**.

### RT-qPCR for transcription expression analysis

Total RNA was extracted from *B. burgdorferi* cell pellets with TRIzol Reagent (Invitrogen; 15596026) according to the manufacturer’s instructions after a 30-minute 3,000 × *g* swinging-bucket centrifugation at room temperature. Residual DNA contamination was removed by a 1-hour DNase I treatment (Thermo Scientific; 18047019) followed by phenol-chloroform extraction (Ambion; AM9720) and standard ammonium acetate/ethanol precipitation (41). RNA samples were used for reverse transcription and quantification of BB_0345, BB_0346, BB_0347, and *flaB* rRNA transcripts by the Luna Universal one-step RT-qPCR kit (NEB, E3005), according to the manufacturer’s instructions, on an Applied Biosystems 7500 Fast real-time PCR system. The primer sets for amplification were BorFlaLeo-R-ok (5’-GCTGGTGTGTTAATTTTTGCAG-3’) + FlaW-sense4 (5’-AGCAACTTACAGACGAAATTAATAG-3’) (42), 0345_F (5’-AAACCCTGAGGGGGTCTTTA-3’) + 0345_R (5’-GGGAAGTCTCTTTTTGCATCC-3’), 0346_F (5’-GACCTCCCCCACTACTACC-3’) + 0346_R (5’-ATAGAGGACATGCAAGCAAC-3’), and 0347_F (5’-ACCAAAAGAAAATGCCTTGC-3’) + 0347_R (5’-CAAGCCTATTTTTGGCGTTT-3’). Transcript levels were validated and normalized against *flaB* endogenous control transcript with fold changes calculated using the comparative CT (2−ΔΔCT) method for quantification.

### Recombinant protein expression

The sequence encoding mature *B. burgdorferi* LolA without a signal peptide was amplified by PCR from *B. burgdorferi* strain B31 genomic DNA (*BB_0346* ORF) using oligonucleotide primers Nde_Q18BblolA_F (5’-GGAATTCCATATGCAAATATCTGCAAATC-3’) and Xho_BblolA_R (5’-CCGCTCGAGATTTTTTTTAATATCAT-3’). This amplicon was NdeI/XhoI restricted and T4 ligated into pET29b(+) expression vector, resulting in pET29b:mlolA-his. After confirmatory sequencing of the insert and flanking regions, the plasmid was transformed into *E. coli* BL21(DE3)pLysS chemically competent cells (Invitrogen, C606003) per manufacture instructions. A single transformant colony on LB agar containing 30 µg/mL kanamycin and 34 μg/mL chloramphenicol was grown overnight in 5mL of selective LB broth with shaking at 37 °C. This starter culture was diluted 1:50 in 1000 mL of selective LB broth and cultured with shaking at 37 °C until an OD_600_ of 0.5. At this point, recombinant protein expression was induced by the addition of IPTG (1 mM final), and incubation was continued for 3 hours. Next, the cells were harvested by centrifugation in a Sorvall RC 6 Plus for 20 min at 5,000 rpm (∼4,400 x g). Cell pellets were resuspended to a total volume of 200 mL in cobalt column binding buffer (50mM NaPO_4_, 300mM NaCl, pH7.4) and frozen at -80°C overnight. After thawing on ice, the resuspended cells were finally lysed with 2 passages through a French press, and the resulting clarified lysate was centrifuged at 10,000 x g for 30 min to collect supernatant containing the soluble protein of interest.

### IMAC and ion exchange FPLC

To purify mature hexahistidine-tagged BB0346 (LolA_Bb_), the soluble lysate was passed through a 0.45 µm sterile filter for loading onto a binding buffer equilibrated HiTrap TALON column via the ÄKTA start protein purification system. After the sample was applied to the column, 15 column volumes (CVs) of 96.7% binding buffer and 3.3% elution buffer (50mM NaPO_4_, 300mM NaCl, 150mM imidazole, pH7.4) were used to wash out unbound protein. Linear gradient elution began with 3.3% elution buffer and ended with 100% elution buffer to obtain an ultraviolet eluent peak between 27% and 63% elution buffer. The collected peak fractions were placed into a 3000 NMWL centrifugal filter unit and concentrated until approximately 1mL of purified sample remained. For subsequent cation exchange chromatography, concentrated affinity column eluent was mixed with 14mL of exchange buffer (50mM MES, pH 5.6), and 4mL of the resulting sample was loaded onto an exchange buffer equilibrated HiTrap SP HP column. After the sample was applied to the column, 15 CVs of exchange buffer were used to wash out any unbound protein. Linear gradient elution proceeded from 0% to 100% ionic elution buffer (50mM MES, 1M NaCl, pH 5.6) with an ultraviolet eluent peak between 38% and 50% ionic elution buffer. A final round of TALON column purification was performed to help eliminate any remaining impurities and to concentrate the protein.

### Crystallization and data collection

Purified LolA_Bb_ containing a C-terminal His-tag was concentrated to 10 mg/mL in 450 mM NaCl, 50 mM MES pH 5.6, 10% glycerol for crystallization screening. All crystallization experiments were set up using an NT8 drop setting robot (Formulatrix Inc.) and UVXPO MRC (Molecular Dimensions) sitting drop vapor diffusion plates at 18 ^°^C. 100 nL protein and 100 nL crystallization solution were dispensed and equilibrated against 50 µL of the crystallization solution. Crystals of LolA_Bb_-PEG were observed after 2 weeks from various conditions in the Berkeley Screen (43) (Rigaku Reagents). A cryoprotectant solution composed of 80% crystallization solution and 20% glycerol was dispensed (2 µL) onto the drop, and crystals were harvested immediately and stored in liquid nitrogen. Crystals used to prepare iodine heavy atom derivatives were obtained from condition F6 (2 M sodium formate, 100 mM HEPES pH 7.5, 5% (w/v) PEG 5000 MME). A solution containing 6.5 µL of crystallant F6, 1.5 µL of 1M potassium iodide (150mM) and 2µL of glycerol was layered onto the drop containing crystals and incubated for 2 minutes. Samples were harvested directly and stored in liquid nitrogen. Crystals of LolA_Bb_-SA were obtained from Berkeley condition E4 (1500 mM Ammonium sulfate, 100 mM Hepes pH 7.5) and samples were cryoprotected with 2.5M lithium sulfate layered onto the drop. X-ray diffraction data for LolA_Bb_-PEG were collected at the Advanced Photon Source beamline 17-ID (IMCA-CAT) and LolA_Bb_-SA crystals were examined a the National Synchrotron Light Source II (NSLS-II) NYX beamline 19-ID. Native data for LolA_Bb_-PEG were collected using crystals from condition F4 at λ=1.0000 Å. Data for the crystals soaked in the presence of potassium iodide from condition F6 were collected at λ=1.7463 Å.

### Structure solution and refinement

Intensities were integrated using XDS (44, 45) via Autoproc (46) and the Laue class analysis and data scaling were performed with Aimless(47). Structure solution was conducted by SAD phasing with Crank2 (48) using the Shelx (49), Refmac (50), Solomon (51), Parrot (52) and Buccaneer (53) pipeline via the CCP4 (54) interface. Six iodide sites were located with occupancies greater than 0.20 and phasing/density modification resulted in a mean figure of merit of 0.58 in the space group *P*4_1_32. Subsequent model building utilizing density modification and phased refinement yielded *R*/*R*_free_ = 0.262/0.294 for the initial model. This model was used for molecular replacement with Phaser (55) against the higher resolution native data set and the top solution was obtained for a single molecule of LolA_Bb_ in the asymmetric unit in the space group *P*4_1_32 (RFZ=3.7, TFZ=82.9, LLG=8408). Additional refinement and manual model building were conducted with Phenix and Coot (56) respectively. Disordered side chains were truncated to the point for which electron density could be observed. Structure validation was conducted with Molprobity (57) and figures were prepared using the CCP4MG package (58). Crystallographic data are provided in **Supplemental Table S1**.

### Growth curves

Cultures of the *B. burgdorferi* 297 conditional knockout strain were inoculated at 1 × 10^5^ cells/mL in selective BSK-II media with various IPTG concentrations. At 24-hour intervals over a 4-day period, cell density was assessed by direct counting of bacterial cultures diluted 2- to 100-fold in PBS using a Petroff Hauser counting chamber under a phase-contrast microscope (Nikon Eclipse E400).

### PK recovery assay for nascent surface lipoprotein transport analysis

After 24 hours with or without 0.1mM IPTG induction of BB0346/LolA_Bb_ expression, 250 mL of late-log spirochetes were harvested at 3,000 x *g* for 30 minutes. The resulting cell pellets were gently washed with 40 mL of Dulbecco′s phosphate buffered saline with 5mM MgSO_4_ (dPBS+Mg) to remove culture medium BSA and re-pelleted at 3,000 x *g* for 10 minutes. Next, the washed cell pellets were gently resuspended in 12.5 mL of dPBS+Mg, and the resuspensions were split into two separate 6 mL aliquots. To one aliquot from each condition (cells grown with or without IPTG), 250 µL of milliQ water was added as a control. To the second aliquot from each condition, 250 µL of 5 mg/mL proteinase K (PK) in milliQ water was added for a final concentration of 200 µg/mL PK. Samples were nutated at room temperature for 45 minutes on the lowest possible speed (approximately 5 rpm). After the incubation, each sample was added to separate 50 mL conical tubes containing 30 mL of dPBS with 5% BSA. The tubes were inverted for mixing, and cells were re-pelleted at 3,000 x *g* for 10 minutes. Resulting cell pellets were gently resuspended in 45mL of selective BSK-II media with or without 0.1mM IPTG, matching the condition they were previously grown in. After a 4-hour recovery incubation at 34°C with 5% CO_2_, the previous steps were repeated for a second round of PK treatment. Specifically, each 45mL culture was harvested at 3,000 x *g* for 30 minutes. The resulting cell pellets were gently washed with 20 mL of BSA-stripping buffer and re-pelleted at 3,000 x *g* for 10 minutes. Next, the washed cell pellets were gently resuspended in 3.125 mL of dPBS+Mg, and the resuspensions were split into two separate 1.5 mL aliquots. To one aliquot from each condition (cells grown with or without IPTG, with or without an initial round of shaving), 62.5 µL of milliQ water was added as a control. To the second aliquot from each condition, 62.5 µL of 5 mg/mL proteinase K (PK) in milliQ water was added for a final concentration of 200 µg/mL PK. Samples were nutated at room temperature for 45 minutes on the lowest possible speed. To each aliquot of cells, 8.25 µL of 1M PMSF protease inhibitor was added for a final concentration of 5 mM PMSF. Finally, the cells were pelleted at 16,000 x *g* for 10 minutes and resuspended in 1x SDS Sample Buffer for analysis by Coomassie-stained SDS-PAGE and Western immunoblots.

### Hypotonic citrate fractionation for outer membrane analysis

OMVs were isolated as previously described (30). Briefly, cultures were harvested and washed with dPBS containing 0.1% (wt/vol) BSA. Cells were then resuspended in 25 mM sodium citrate (pH 3.2) containing 0.1% (wt/vol) BSA. Cell suspensions were shaken for 2 hours at room temperature in a New Brunswick C24 incubator at 250 rpm to release OMVs. After this agitation period, the cell suspensions were harvested, resuspended in citrate buffer containing BSA, and loaded onto a discontinuous 56%-42%-25% (wt/wt) sucrose gradients prepared in Beckman UltraClear tubes. The gradients were centrifuged at 120,000 × g for 18 hours at 4°C in a Beckman Coulter XPN-80 ultracentrifuge using an SW 32 Ti rotor, and the resulting upper outer membrane vesicle (OMV) bands and lower protoplasmic cylinder (PC) bands were extracted by needle aspiration. Fractions were diluted in cold dPBS, re-pelleted separately, and then resuspended in dPBS containing 1 mM PMSF. A portion of the resuspended fractions was used to prepare SDS-PAGEs and western immunoblots, and the remainder was stored at −80°C for later analysis.

### Genomic sequencing of *B. burgdorferi* 297

Hybrid Oxford Nanopore Technology (ONT) and Illumina short read sequencing was performed by SeqCenter (Pittsburgh, PA) on genomic DNA isolated from *B. burgdorferi* strain 297 with a Quick-DNA Miniprep Plus Kit (Zymogen, D4068). Specifically, Illumina sequencing libraries were prepared using the tagmentation-based and PCR-based Illumina DNA Prep kit and custom IDT 10bp unique dual indices (UDI) with a target insert size of 320 bp. No additional DNA fragmentation or size selection steps were performed. Illumina sequencing was performed on an Illumina NovaSeq 6000 sequencer in one or more multiplexed shared-flow-cell runs, producing 2 x 151bp paired-end reads. Demultiplexing, quality control and adapter trimming was performed with bcl-convert1 (v4.1.5). Nanopore sequencing was performed on an Oxford Nanopore a MinION Mk1B sequencer or a GridION sequencer using R10.4.1 flow cells in one or more multiplexed shared-flow-cell runs. Run design utilized the 400bps sequencing mode with a minimum read length of 200 bp. Adaptive sampling was not enabled. Guppy1 (v6.4.6) was used for super-accurate basecalling (SUP), demultiplexing, and adapter removal.

### Assembly and annotation of *B. burgdorferi* strain 297 genomic sequencing data

ONT and Illumina reads were *de novo* assembled using a combination of Trycycler long-read assembly, Medaka long-read polishing, and Polypolish short-read polishing, as described by Wick and colleagues (59). The main linear chromosome (965,079 bp) and circular plasmid cp26 (26,514 bp) were identified from the resulting assembled contigs, with cp26 having 100% sequence identity to a previously deposited strain 297 cp26 sequence (Genbank accession number CP002268.1) (60).

11 kb and 33 kb reverse complement (i.e. inverted repeat) sequences flanking both ends of the large linear chromosome were manually removed from the assembly. These sequencing artifacts arose from the covalently closed hairpin telomeres of linear genomic elements in *B. burgdorferi* (61). Because the smaller plasmid elements proved challenging to assemble *de novo* with high confidence, a hybrid FASTA file was generated, combining the new linear chromosome sequence with all 20 strain 297 plasmid sequences previously assembled by Schutzer and colleagues (60). This FASTA file was run through the National Center for Biotechnology Information (NCBI) Prokaryotic Genome Annotation Pipeline (PGAP), and arbitrary locus tags (*e.g.* pgaptmp_000001) were replaced with known *B. burgdorferi* strain B31 locus tags (*e.g.* BB_0001) in the resulting GenBank file by local BLASTP analysis of the *B. burgdorferi* 297 proteome against the proteome of *B. burgdorferi* B31 (Assembly ASM868v2). The PGAP-annotated assembly of the complete *B. burgdorferi* 297 genome, including the here assembled linear chromosome prior to flanking sequence removal, together with the BLAST-based ORF annotation file, is provided in the **Supplemental materials**.

### Analysis of cells fractions by quantitative label free mass spectrometry

50 µg of each cell fraction was precipitated by trichloroacetic acid (TCA)/acetone as described (62) for analysis by multidimensional protein identification technology (MudPIT) mass spectrometry (63). Briefly, peptides eluted at every step of a 10-step LC/MS process were searched against a *Borrelia burgdorferi* 297 protein sequence database using the ProLuCID search engine. The result files from the ProLuCID search engine were processed with DTASelect (v 1.9) to assemble peptide level information into protein level information. In-house software, swallow and sandmartin (v 0.0.1), worked with DTASelect to select Peptide Spectrum Matches such that the FDRs at the peptide and protein levels were less than 1%. Peptides and proteins detected in the 12 samples were compared using CONTRAST. Proteins that were subsets of others were removed using the parsimony option in DTASelect after merging all runs. Proteins that were identified by the same set of peptides (including at least one peptide unique to such protein group to distinguish between isoforms) were grouped together, and one accession number was arbitrarily considered as representative of each protein group. In-house quantitative software, NSAF7 (v 0.0.1), was used to create a quantitative Contrast Report on all detected peptides and non-redundant proteins identified across the different runs. These data were analyzed in R with filtering for proteins detected in at least 2 of the 3 biological replicates for each fraction. The standard error for fraction ratios was calculated with propagation of uncertainty. All proteins with significant OMV and PC abundance changes identified by this analysis are listed in Supplemental Tables S2 and S3, respectively.

### SDS-PAGE and immunoblotting for protein expression analysis

*B. burgdorferi* cells were harvested by centrifugation in a swinging bucket at 3,000 × *g* for 30 min at room temperature. Harvested cells were washed twice with dPBS+Mg and resuspended in standard 1× SDS sample buffer (41). Whole-cell lysates were resolved by 12% or 18% SDS-PAGE and visualized by Coomassie staining (Fisher; BP3620-1). For immunoblotting, proteins were electrophoretically transferred overnight at 4°C to 0.1 µm nitrocellulose blotting membrane (GE, 10600000) using a Bio-Rad Mini Trans-Blot apparatus at 30 V with prechilled transfer buffer (25 mM Tris, 200 mM glycine, and 20% methanol). Membranes were blocked for 30 min in TBST buffer (25 mM Tris, 150 mM NaCl, 0.05% Tween 20, pH 7.2) with 5% dry milk before incubating on a rocker overnight at 4°C with either mouse anti-FLAG (1:5,000 dilutiion; Thermo Scientific, MA191878), rat anti-BB0346-HIS (1:5,000 dilution), rat anti-FlaB (1:4,000 dilution; reference (64), mouse anti-OspA (1:10,000 dilution; H5332; reference (65)), rat anti-OppAIV (1:5,000 dilution, a gift from M. Caimano, University of Connecticut Health Center), mouse anti-P66 (1:500 dilution; H1337; reference (66)), or mouse anti-Lp6.6 (1:10; reference (7)) primary antibodies. After three 10-minute washes with TBST, the blots were incubated at room temperature on a rocker for 1 h with secondary anti-mouse IgG-HRP (Sigma; A4416) or anti-rat IgG-HRP (Thermo Scientific; 31470) antibody. After three additional 10-minute TBST washes, membranes were allowed to react with Super Signal West Femto (Thermo Scientific; 34096) substrate per manufacturer instructions. Chemiluminescence was detected with automatic exposure settings on an Amersham ImageQuant™ 800 (Cytiva).

### Accession codes

*B. burgdorferi* LolA coordinates and structure factors have been deposited to the Worldwide Protein Databank (wwPDB) with the accession codes 7TPM (PEG bound) and 8T5T (steric acid bound). Assembled genome sequences for *B. burgdorferi* 297 are available in the GenBank database under accession numbers CP152378 (chromosome) and CP152379 (cp26). Raw MudPIT label free quantitative mass spectrometry data are available in the MassIVE database under accession number MSV000095380.

## ACKNOWLEDGEMENTS

We thank Jing Liu for the initial cloning of the *B. burgdorferi lolA* sequence, Hajime Tokuda for sharing *E. coli* strains, and Jessica Kueker for the preliminary complementation experiments. Work at KUMC was supported in part by NIH/NIAID Grants R21AI144624 and R21AI178835 and KU Endowment/Bryan Lynch Family Foundation (to W.R.Z.). Work at UAMS was supported by NIH/NIAID grants R01AI087678 and R21AI119532 (to J.S.B.), the UAMS Center for Microbial Pathogenesis and Host Inflammatory Responses (supported by NIH/NIGMS grant P20GM103625 to J.S.B.), and the UAMS IMSD program (NIGMS R25GM083247 to D.E.G.). Use of the KU Protein Structure Laboratory was supported by NIH/NIGMS grant P30 GM110761. This project has also been in part supported by Federal funds from NIH/NIAID/DHHS Contract No. 75N93022C00036. Use of the IMCA-CAT beamline 17-ID at the Advanced Photon Source was supported by the companies of the Industrial Macromolecular Crystallography Association through a contract with Hauptman-Woodward Medical Research Institute. Use of the Advanced Photon Source was supported by the U.S. DOE, Office of Science, Office of Basic Energy Sciences, under Contract No. DE-AC02-06CH11357. This research used resources the NYX beamline 19-ID, supported by the New York Structural Biology Center, at the National Synchrotron Light Source II, a U.S. DOE Office of Science User Facility operated for the DOE Office of Science by Brookhaven National Laboratory under Contract No. DE-SC0012704. The NYX detector instrumentation was supported by grant S10OD030394 through the Office of the Director of the NIH. S.K.S. and L.F. were supported by the Stowers Institute.

